# The future is faeces: Using faecal genomic sequencing to understand dietary choices of an endangered arboreal marsupial

**DOI:** 10.64898/2025.12.10.693363

**Authors:** Mikaela J. Bratovic, Celine H. Frère, Nicola Jackson, Linda Moss, Michaela D. J. Blyton, Dominique A. Potvin

## Abstract

**1.** Understanding critical components of habitat use, such as dietary preferences, is crucial for creating and implementing effective conservation plans. This becomes challenging when species exhibit cryptic behaviours, such as arboreality and nocturnality, as seen in the endangered greater glider *(Petauroides spp)*. Existing methods for determining greater glider diet are inadequate, time consuming and prone to human error. Furthermore, current literature lacks specificity, with most research stating simply that greater gliders are dietary eucalypt specialists, without providing additional insight into which species are eaten. **2.** Here, we tested a non-invasive technique using targeted sequencing of species-specific single nucleotide polymorphisms from greater glider faecal samples to identify dietary components across multiple locations in Southeast Queensland. **3.** Sequencing identified 17 plant species present in the faecal samples, including *Angophora, Acacia* and *Casuarina spp.* that were previously unknown to be feed tree taxa, profoundly increasing our understanding of the habitat requirements of this endangered marsupial. Four of these identified species were out of range of their natural occurrence, suggesting limitations to the accuracy of this technique. **4.** This study demonstrates the value of using genomic sequencing for analysing diets of arboreal mammals and makes recommendations for improving the accuracy of this methodology for future studies. Our findings highlight key tree species to be considered important for future greater glider conservation plans and raises the question whether there are local drivers behind the differences in dietary choices across the landscape. Such information will be critical for greater glider conservation, particularly when management planning across multiple habitats.

## Introduction

A sound understanding of the ecology of species threatened with extinction (hereon “threatened”) is vital for ensuring effective conservation and management plans for species recovery (Castle et al., 2020; Lindenmayer et al., 2011; Martin et al., 2022). Rare or “uncommon” threatened species, based on geographic range, scarcity of population numbers and infrequent sightings in the wild, are a key focus for conservation ecology, but collecting data on these species is a challenge (Martin et al., 2022). This difficulty is amplified when the threatened species exhibit cryptic behaviours including nocturnality and arborealism (Martin et al., 2022; Neilson et al., 2013). Previous studies suggest that without robust, empirical data on threatened species’ resource preferences, conservation decisions can unintentionally become disadvantageous as they are being based on outdated, sparse, or inaccurate information (Bogert, 1994; Bryant et al., 2016; Joly et al., 2010; Martin et al., 2022).

Understanding dietary components of threatened species’ resource preferences is one crucial area contributing to their conservation both *in situ* and *ex situ* (Birnie-Gauvin *et al*., 2017; Castle *et al*., 2020; Oftedal & Allen, 1996). This involves exploring feeding preferences and avoidances, quantifying nutritional needs, recording foraging patterns and understanding digestive physiology (Birnie-Gauvin et al., 2017; Jordan, 2006). Identifying feeding preferences of folivores, frugivores and granivores is of particular importance to *in-situ* conservation as it highlights the target tree species to be used for effective habitat restoration and protection (Kowalczyk *et al*., 2019; Neilson *et al*., 2013; Zhong *et al*., 2023). This can act as a critical framework for implementing legislation to aid in threatened species recovery. Expectedly, analysing the diet of cryptic species is challenging, but the field of dietary analysis is rapidly evolving regarding efficiency and accuracy (Blyton *et al*., 2023; Jones & Krockenberger, 2007; Kowalczyk *et al*., 2019; Nielsen *et al*., 2018).

Historically, diet analysis has been performed using techniques that rely heavily on the observer’s judgement and base knowledge, often being susceptible to bias (Gilby et al., 2010; Holechek et al., 1982; Nielsen et al., 2018). Direct observational surveys such as continuous focal follows, which involves recording the behaviour of an individual for a predetermined period of time, or scan sampling, which involves recording the behaviour of an individual at predetermined intervals, are commonly used observational methods to record what a subject is eating at a given point in time (Gilby et al., 2010). However, it is often difficult for researchers to identify food species using direct observational methods, particularly in dense habitats, and observations may not include timeframes long enough to capture the full breadth of diet items consumed (Matthews *et al*., 2020; Parker & Bernard, 2006). For example, a comparative study on methods of dietary analysis in chimpanzees (*Pan troglodytes schweinfurthii*) found that scan sampling intervals completely omitted the consumption of insects, which were otherwise recorded in the chimpanzee’s diets using alternate sampling methods (Matthews *et al*., 2020). Alternatively, macroscopic inspection of gut contents allows for a retrospective approach to identify what has recently been ingested (Holechek et al., 1982; Jordan, 2006; Matthews et al., 2020) and provides a more precise method for determining the exact contents consumed. However, this also only provides a snapshot of diet at one time, and the invasiveness of the procedure limits its applicability in many circumstances (Holechek et al., 1982; Wilson *et al*., 1977). Furthermore, it can be challenging to accurately identify partially digested foods, particularly plant species, due to their differential digestion rates (Holechek et al., 1982).

Developments in the field of genetics have allowed the introduction of DNA extractions for accurate dietary analysis. Genetic extraction from faecal samples provides a non-invasive method to identify components found within the faeces, indicating what species the animal has ingested (Liu et al., 2021; Moritz & Cicero, 2004; Pompanon et al., 2012). Genetic dietary analysis has generally been performed using metabarcoding, which effectively increased accuracy compared to traditional visual observation methods (Castle et al., 2020; Kowalczyk et al., 2019; Quéméré et al., 2013). However, the DNA fragments amplified in metabarcoding are often unable to provide enough resolution to classify consumed taxa to species level (Liu et al., 2021; Pompanon et al., 2012). Studies suggest that using metabarcoding to analyse herbivorous species’ diets presents a greater challenge than carnivores’, due to this lack of resolution when delineating plant species (Blyton et al., 2023; Bradley et al., 2007; Pompanon et al., 2012). Blyton *et al*. (2023) highlighted the inability to use metabarcoding on animals consuming eucalypts, due to frequent hybridisation and other genetic similarities between eucalypt taxa. This has prompted the use of a new technique for genetic dietary analysis using next generation sequencing of single nucleotide polymorphisms (SNPs) (Blyton et al., 2023).

Like metabarcoding, SNP-based analysis identifies species-specific markers, however, instead of targeting entire DNA segments, they detect variations in single base pairs, offering greater taxonomic resolution (Blyton et al., 2023; Komar, 2009). Previous studies have confirmed the ability to use SNPs to identify most eucalypts to species level (Dasgupta et al., 2015; Schultz et al., 2018; Yuskianti et al., 2011), and Blyton et al. (2023) was the first study to successfully use SNPs to conduct a comprehensive dietary analysis on a herbivorous species. In their study, they used the DarTseq platform (Diversity Arrays Technologies) to sequence a reduced fraction of the genomes of potential food tree species (Jaccoud et al., 2001; Kilian et al., 2012). This sequence data was then used to identify SNPs that were species-specific within the set of reference species. Oligo’s were then designed to amplify selected species-specific SNPs (Diet Oligo Panel) from scat DNA that were then sequenced via the DarTag platform to identify koala *(Phascolarctos cinereus)* diet composition (Blyton et al., 2023). The effectiveness of this method is yet to be verified on other herbivorous species, but its application appears transferrable to those with diets potentially similar to koalas.

Greater gliders (*Petauroides spp.)* are one taxon that could benefit greatly from using SNP-based dietary analysis. These threatened marsupials are endemic to the eastern mainland of Australia where habitat fragmentation, deforestation, bushfires and inappropriate fire regimes are causing a rapid reduction in numbers with a population decline of more than 50% over a 21-year period (Australian Government, 2022; Lindenmayer et al., 2022; Nelson et al., 2018; Wagner et al., 2020). Until recently, greater gliders were classified under the sole species of *Petauroides volans,* with the Australian Government recognising two subspecies (*P. v. minor* and *P. v. volans*) for conservation purposes (Australian Government, 2016). More recent studies based on morphological differences (Jackson & Groves, 2015) and genetic analysis (McGregor et al., 2020) have recognised three distinct species; *Petauroides volans* (southern), *P. minor* (northern), and *P. armillatus* (central). Although a rough understanding of the distributions of each species has been described, the lack of sampling and the addition of hybridisation has made the exact zoning unclear (Australian Government, 2022; Jackson & Groves, 2015; McGregor et al., 2020).

Regardless of species delineations, little is known about greater gliders’ dietary needs or preferences. Accurate identification of dietary components would allow us to ensure that access to these species is prioritised in habitat protection decisions and reforestation. Current conservation plans and research have a strong focus on landscape-based management, especially patch size, movement corridors and the presence of tree hollows (Hofman et al., 2022; Norman & Mackey, 2023). Although these factors are crucial for quality greater glider habitat, having a comprehensive knowledge of their diet to couple with these landscape features is paramount to optimise conservation efforts. It is currently accepted in the literature that greater gliders are strictly folivores, feeding exclusively on eucalypts, but this description remains vague. While captive feeding experiments have confirmed that greater gliders will consume foliage from the *Eucalyptus* and *Symphyomyrtus* subgenera, with some studies focussing to species level (commonly *Eucalyptus radiata),* this does not inform whether gliders prefer these species in the wild or are just consuming what is provided to them in captivity (Foley et al., 1987; Jensen et al., 2014; Youngentob et al., 2011). All in situ research on greater glider diet has used only visual observation methods of data collection, including either transect spotlighting or radiotracking with spotlighting. Some studies noted the limitation that greater gliders altered their behaviours, often freezing when a spotlight was shone upon them, meaning data collected was not completely indicative of natural behaviours (Kavanagh & Lambert, 1990; Kehl & borsboom, 1984). The transition towards genetic-based diet techniques presents the opportunity to analyse the diet of greater gliders with reduced bias and increased accuracy.

Here, we sought to determine the diet of greater gliders through the use of next-generation sequencing techniques, using a pre-existing QLD SNP Diet Oligo DArTag Panel. This panel was developed as part of a Queensland Government sustainability action grant and has been used to characterise the diets of koalas from over 400 faecal samples from Queensland. Methodology for the panel creation and the markers included in the panel can be found on the project website (Moore & Blyton, 2025). Given the previously recorded similarities between koala and greater glider food preferences (Jensen et al., 2015; Moore et al., 2004), the panel was likely to be effective for identifying greater glider diet. We predicted that greater gliders would show an affinity to a small number of feed tree species consistent across all sampling locations due to previously being described as picky, specialist feeders (Comport et al., 1996; Kavanagh, 1984; Kehl & borsboom, 1984). We substantiate the success of using this non-invasive technique to investigate the diet of a cryptic mammalian species, whilst highlighting shortcomings of the methodology and suggesting improvements to optimise the outcomes for future studies.

## Materials and Methods

### Study Area and Field Methods

The study was conducted within two local government areas in southeast Queensland, Australia, where glider populations are suspected or have been confirmed, and which are under consideration for protection and management; the City of Moreton Bay, and Gympie (Figure 1). The City of Moreton Bay (27.0946° S, 152.9206° E) was the primary study area, and all sampling occurred on council-owned land (Figure 1; Supplementary Table 1). One sample location was included from the Gympie Region (26.03486° S, 152.51390° E) on a private property in Sexton with known greater glider activity. Fieldwork was conducted across two periods. The initial sampling occurred over six days in Autumn 2024 (March – May) and a secondary bout of fieldwork was completed over six days in Spring 2024 (September). Sample locations within the City of Moreton Bay were determined by assessing historical maps and identifying potential greater glider habitat based on known preferred characteristics including the presence of old growth trees and remnant vegetation patch size.

**Figure 1.**
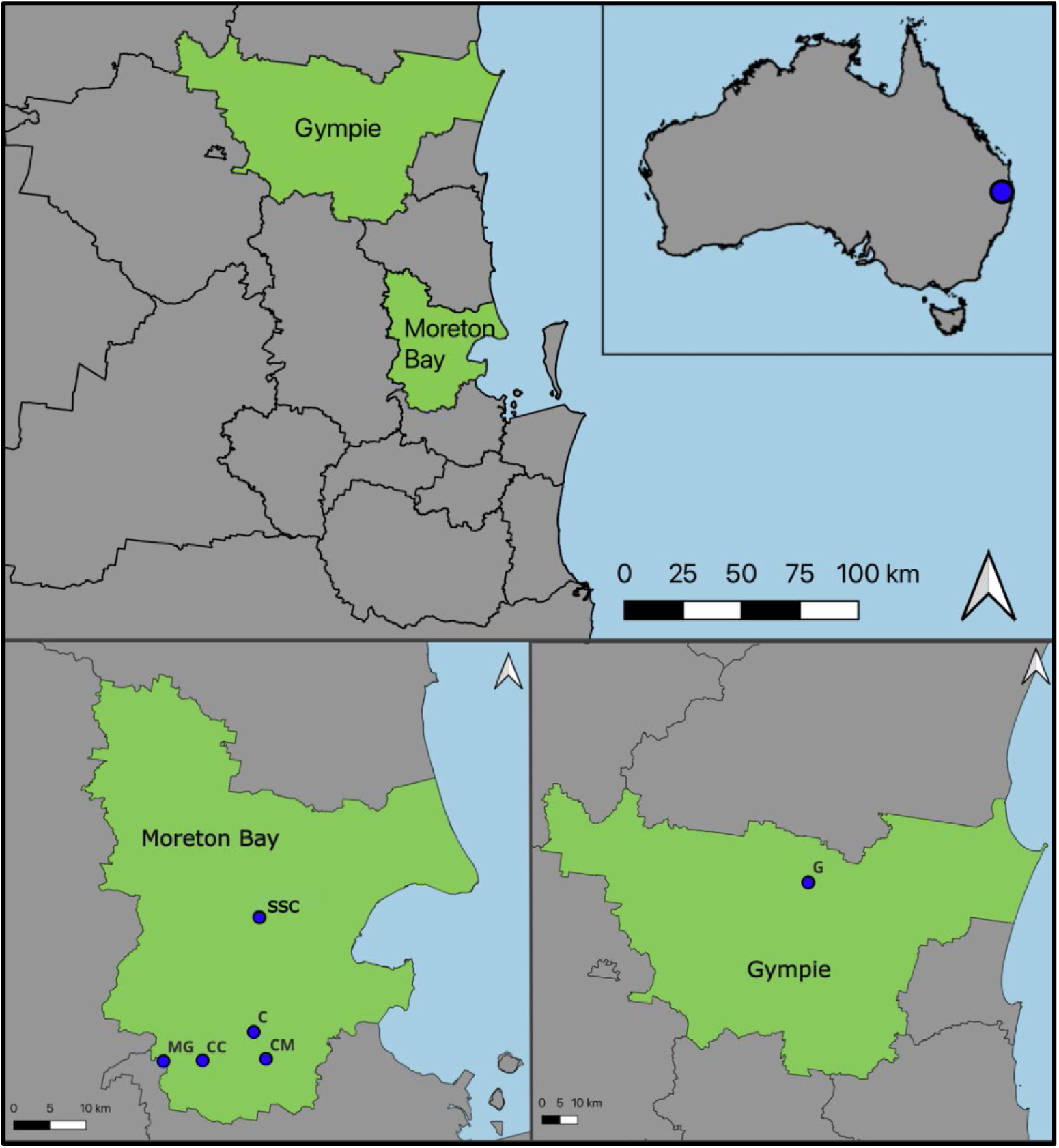
Location of the two local government areas used as study sites: The City of Moreton Bay and Gympie Regional Council. Blue dots within the local government areas indicate individual sites where greater glider faecal samples were collected. SSC – Sheep Station Creek, C – Cashmere, CM – Clear Mountain, CC – Cedar Creek, MG – Mount Glorious, G – Gympie.

Within the City of Moreton Bay, a detection dog and her handler (Nicky Wright - Animal Behaviour Consultant - Morekos) were employed to locate greater glider faecal samples. The dog, who had been trained to specifically detect greater glider faeces, was directed by her handler to scout along a transect for any samples, with the freedom to venture away from the transect line when following a scent cone. Once the dog indicated to a sample spot, the handler marked the location with an identifying flag. The area in a five cm radius surrounding the flag was then searched visually by researchers. If no faeces were identified by researchers within 30 seconds, the flagged location was abandoned. If faecal samples were identified, they were collected into a labelled falcon tube and maintained on ice before storing samples at −20°C. The GPS location was recorded each time a sample was collected.

Within the Gympie Region, one known greater glider habitat was sampled visually for faecal samples. This was conducted by searching around the base of trees that had nest boxes previously deployed for gliders by the property owner, natural hollows or that appeared large enough to be utilised by greater gliders in any capacity. Any suspected faecal samples were collected in labelled test tubes and stored on ice until placed into a −20°C freezer. Sampling in the Gympie Region occurred over one day in August 2024.

### Host Species Identification

First, host DNA was extracted from the outer layer of the faecal samples, either by rolling the pellet onto the surface of paraffin gauze or using a flocked swab dipped in phosphate buffered saline (PBS) wiped over the surface. This aimed to sample the epithelial cells that are attached to the outer surface of the scat as it moves through the digestive tract, similar to established methods used in koalas (Schultz et al., 2018). A Qiagen DNeasy Blood & Tissue Kit was used to extract the DNA according to the manufacturer’s instructions with the following additional minor modifications. Prior to commencing the kit instruction, 500mL of PBS was added to the tubes containing the gauze. This was heated at 57°C for 10 minutes then centrifuged at 18000 rpm for 5 minutes to melt and separate the paraffin before continuing with the Qiagen protocol. Changes to the original protocol included a 1hr incubation at 56°C to lyse cells, and a final elution volume of 100µL of AE buffer. After extraction, the DNA was stored at −20°C for downstream testing and analysis.

Primers targeting cytochrome-*b* on the mitochondrial genome were used for identifying the host species for the initial Moreton Bay samples and the Gympie samples. The L14724 and H15149 universal cytochrome-*b* mammalian primers from Irwin et al. (1991) were redesigned by an external laboratory to target greater gliders and possums (F. Hogan & F. Wedrowicz, unpublished data). The resulting primers used to create the DNA fragments were cytB-F (5’ – CCTATGGCATGAAAAACCATTGTTG – 3’) and cytB-R (5’ – CCTCAGAAAGATATTTGGCTCATGG – 3’). PCR was conducted using 10µL AmpliTaq Gold 360 (2X) master mix (Thermo Fisher Scientific), 0.5µL of both the forward and reverse primers at 10µM concentration, 7µL of PCR grade water and 2µL of template DNA. The thermal cycling profile was 98°C for 1 min, followed by 35 cycles of 98°C for 15 s, 57°C for 15 s, and 72°C for 30 s, with a final extension of 72°C for 1 min. Non-template controls (PCR free water) and ringtail possum (*Pseudocheirus peregrinus*) DNA positive controls were also included. All samples were run on an electrophoresis gel with 2% agarose and SYBR Safe gel stain (Thermo Fisher Scientific) to confirm amplification of host DNA.

All samples exhibiting DNA amplification on the gel were sent for Sanger sequencing at the Australian Genome Research Facility, St Lucia, Australia. The resulting forward and reverse sequences were aligned in Geneious Prime (v 2024.0.5) using the de novo assembly function. Reference genomes from tissue samples for greater gliders (*Petauroides volans)* and closely related gliding and possum species found in the study area *(Pseudocheirus peregrinus, Trichosurus caninus, Trichosurus vulpecula, Petaurus breviceps)* were used to cluster the sequences and identify the host species of each sample.

A greater glider specific primer developed by co-authors targeting the 16S region of the mitochondrial genome was used to identify the host DNA of the secondary samples from Moreton Bay (Jackson et al., 2026 forthcoming). For this, nine greater glider mitochondrial genomes spanning all three recognised species were aligned using MAAFT (V7.409). Geneious V2025.03 was used to identify primers specific to all three species, whilst using all available mitochondrial references for Australia mammals as an off-target database. NCBI *blastn* was used to ensure no off-target amplification with other species. Primers resulted in an approximate 153bp amplicon specific to the greater glider species complex. A PCR was conducted with extracted faecal sample DNA using 10µL AmpliTaq Gold 360 (2X) master mix (Thermo Fisher Scientific) and 0.5µL of both the forward and reverse primers at 10µM concentration. The thermal cycling profile was 95°C for 10 min, followed by 40 cycles of 95°C for 15 s, 60°C for 30 s, and 72°C for 15 s, with a final extension of 72°C for 7 min. A 2% agarose gel made with SYBR Safe gel stain (Thermo Fisher Scientific) was used to visualise the PCR product. Non-template controls and a greater glider DNA positive control were also included. All samples indicating positive greater glider DNA were included for dietary analysis. These included six samples from Sheep Station Creek, four from Mount Glorious, two each from Cedar Creek, Gympie and Cashmere, and one sample from Clear Mountain (locations in Figure 1).

### Faecal DNA Extractions

DNA was extracted from the faecal samples confirmed to be greater glider using a high salt CTAB extraction protocol as described by Inglis *et al* (2018). This involved using a sorbitol pre-wash, 3% CTAB extraction buffer, chloroform clean-up and an isopropanol precipitation. Approximately 100mg of each faecal sample was flash frozen in liquid nitrogen then ground into a fine powder using a TissueLyser with a 5mm stainless steel bead. A sorbitol buffer (100 mM Tris-HCl pH 8.0, 0.35 M Sorbitol, 5 mM EDTA pH 8.0, 1% (w/v) polyvinylpyrrolidone, 1% (v/v) 2-mercaptoethanol) was used to wash the samples before they were mixed with the extraction buffer (100 mM Tris-HCl pH 8.0, 3 M NaCl, 3% cetyl trimethylammonium bromide, 20 mM EDTA and 1% (w/v) polyvinylpyrrolidone, 1% (v/v) 2-mercaptoethanol) and incubated at 65°C for 1 hour. One volume chloroform:isoamyl (24:1) alcohol was used to wash the supernatant. Precipitation occurred with the addition of 3 M sodium acetate pH 5.2 (0.1 x volume) and cold isopropanol (0.66 x volume). The tubes were then incubated in a −20°C freezer for one hour. Following incubation, the samples were centrifuged for 10 minutes at 13000 x g. The DNA pellet was washed with 70% ethanol, then dried out for 10 minutes at 65°C. Once dry, the DNA was suspended in 40 µL TE buffer containing 0.1 mg ml-1 DNase-free RNase A and incubated at 37°°C for 30 minutes. The final product was sent to Diversity Arrays Technology P/L, Canberra, Australia (DArT) for single nucleotide polymorphism (SNP) sequencing using the QLD SNP Diet Oligo DArTag Panel. Prior to sequencing, the samples underwent gDNA purification at DArT using ZYMO technology to ensure the highest success of read counts.

### DArTag sequencing

The QLD SNP Diet Oligo DArTag Panel contains 1029 validated markers (and 93 unvalidated markers) for 89 koala candidate food tree species including an extensive selection of eucalypts (*Corymbia, Eucalyptus* and *Angophora*) and select *Melaleuca, Acacia, Casuarina* and *Lophostemon* species (Moore & Blyton, 2023). Although the results using this method are limited to the species in this panel, and it is not likely to include all possible plant sources for greater gliders, it remains the most comprehensive tool available for determining folivore diets in Queensland. The increased resolution provided by SNP sequencing confirms the advantages of this method over metabarcoding, and the increased number of reads from using targeted oligos provides greater accuracy compared to traditional non-targeted next generation sequencing methods (Blyton et al., 2023). Widening the selection of plant species in this panel would provide more comprehensive outcomes for future studies.

Faecal DNA extracts were sequenced on the DArTag platform using the QLD SNP Diet Oligo DArTag Panel to first amplify species-specific SNPs. The raw sequencing reads were then processed using DArT custom bioinformatics pipelines to produce a raw SNP read count by sample table, DArTSoft 7.4 (Diversity Arrays Technology P/L, Canberra, Australia).

### Determining Diet from Faecal DNA

The raw SNP read count data was returned and filtered by the implementation of the following predetermined parameters to remove likely sequencing errors and low frequency contamination. For each target species, sequencing reads were pruned from the dataset if the average number of reads per detected marker was less than three. Additionally, reads were removed if fewer than two SNPs were detected for a target species that had three or more validated markers. To correct for differences in amplification efficiency between markers during the DArTag sequencing, the filtered reads for each marker were divided by the average number of reads returned for that marker from DArTag sequencing of DNA from the target tree species conducted during the marker validation process. The number of scaled reads were then averaged across the successfully amplified markers for each target species. The average scaled read counts for each target species were then converted to relative abundance by dividing by the total average scaled reads across all target species within a sample. Data analysis was completed in Microsoft Excel (v16.92) with the use of R Studio (v2023.06.1+524)(R Core Team, 2023) to combine individual marker sequences into target population level reads and generate plots with the ‘ggplot2’ package (Wickham, 2016).

### Taxonomic inconsistencies

Once a list of plant species found within the faecal samples was produced, it was looked over to identify any differences between the species names on the DArTag Panel and the current taxonomic standards. Any inconsistencies were amended at this point. The panel included subspecies for a select number of plants, however, this was not consistent across all species. In the interest of continuity, reads for subspecies were combined with their parent species, meaning no subspecies were included in this study.

### Cross-referencing SNP sequencing results with local occurrence

A species list of plants known to occur at each sampling location was collated with data from iNaturalist and Atlas of Living Australia (Supplementary Table 2). This was used to cross-reference with the species identified in the greater glider faecal samples by the DarTag sequencing. Any species from the samples that did not align with the list were then investigated to identify the expected distributions and possible reasons as to why they were detected.

### Animal Ethics

Ethics for the detection dog fieldwork and faecal collection in Moreton Bay was covered under Animal Ethics CA 2023/10/1793 (Conservation Detection Dog Service).

## Results

After filtering and refining the SNPs returned from sequencing 24 faecal samples, there was a total of 4489 reads, across 17 faecal samples, matching with plant species on the QLD SNP Diet Oligo DArTag Panel (Supplementary Material 3). Seven (29%) samples failed to return diet associated reads and another 5 (21%) returned fewer than 50 diet associated reads (Figure 2). From the 17 samples, 17 species of plants were identified from the genera *Corymbia, Eucalyptus, Acacia, Angophora* and *Casuarina* as eaten by our sampled greater gliders in Moreton Bay and Gympie (Figure 2; Table 2). Three plant species, *C. citriodora, C. maculata* and *E. microcorys* were identified in faecal samples across multiple sampling locations, whereas the remainder of the identified plant species were unique to their location. It should be noted that although *E. laevopinea* reads were identified in samples, due to insufficient DNA when developing the QLD SNP Diet Oligo DArTag Panel, the markers for the species remain unvalidated, therefore it cannot be included when producing relative abundance calculations.

**Figure 2.**
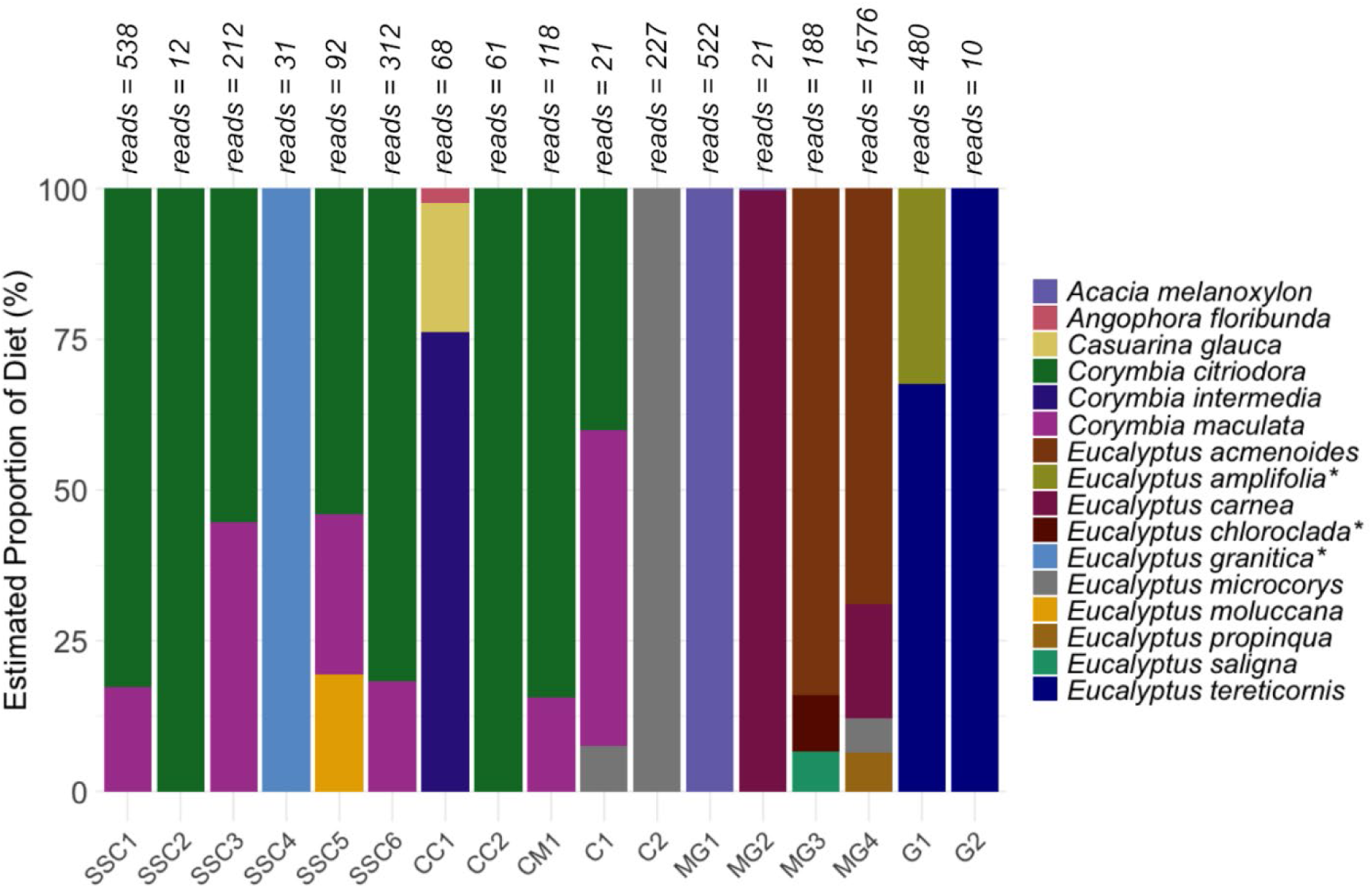
The estimated diet composition of each greater glider faecal sample sequenced with DArTag from Moreton Bay and Gympie. Number of sequenced reads is displayed above each sample bar. *Natural range of species does not overlap with study sites.

**Table 2.**
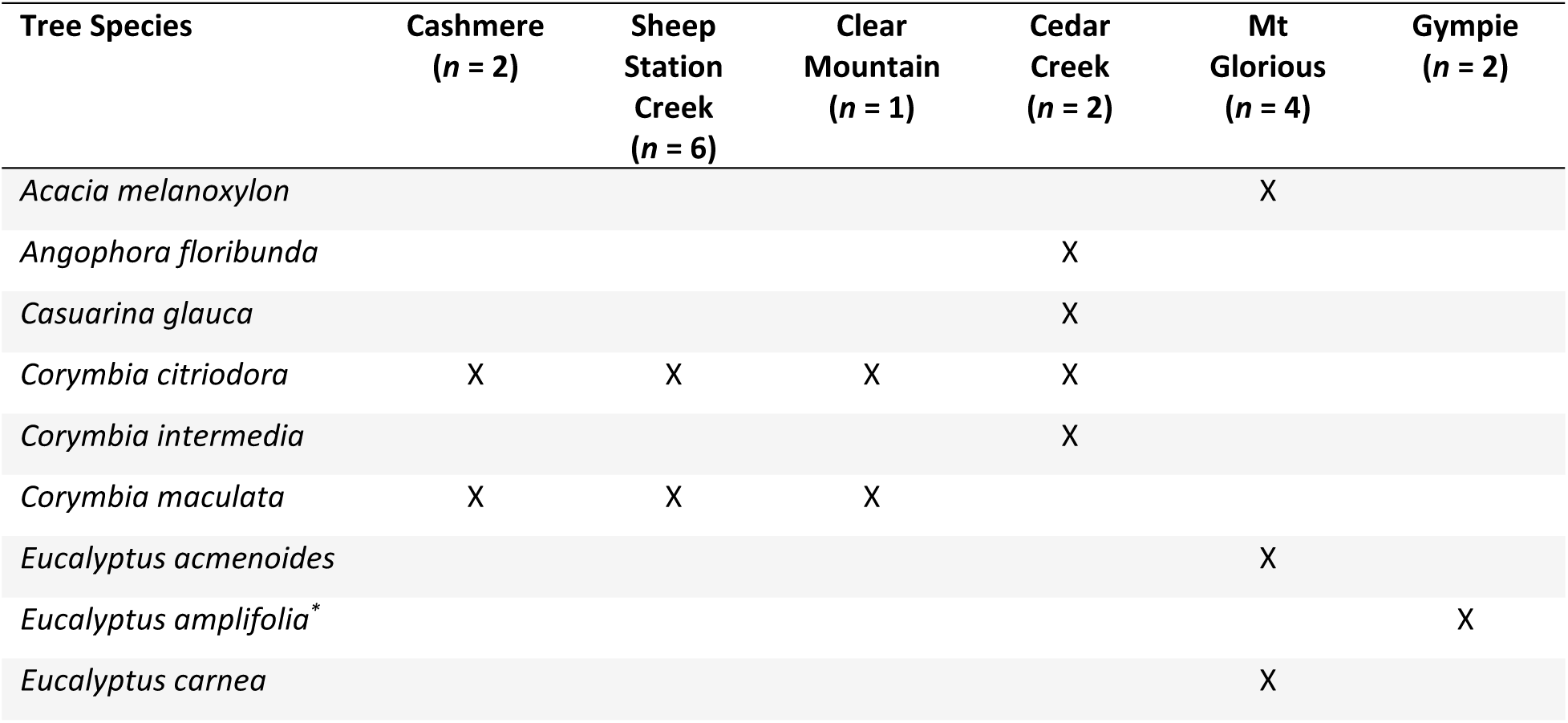
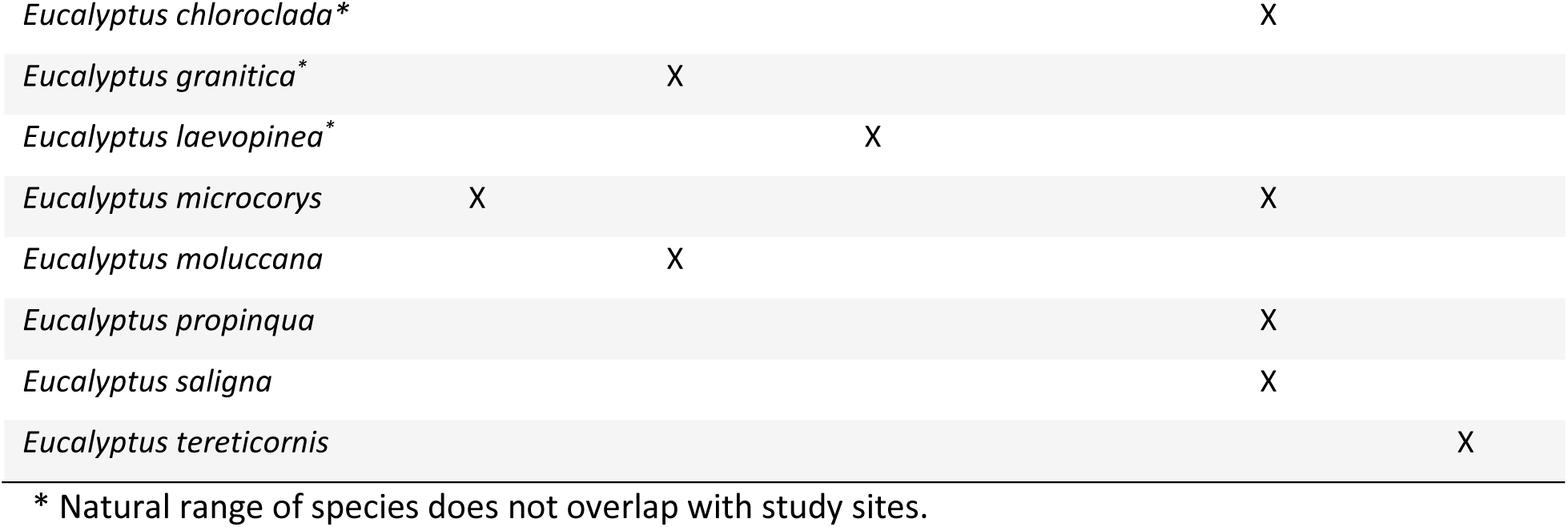
Plant species identified in the faecal samples from DArTag sequencing and their corresponding collection locations across Moreton Bay and Gympie.

From the plant species identified in the greater glider diet, *C. citriodora* and *C. maculata* had the highest frequency of occurrence (Figure 3). Six out of the seventeen plant species were present in more than 10% of the individual faecal samples. A separate six species were discovered to dominate the samples they appeared in, equating for more than 50% of the plant DNA found in their respective samples. *Corymbia citriodora* appeared most frequently, being found in half of the faecal samples (Figure 3).

**Figure 3.**
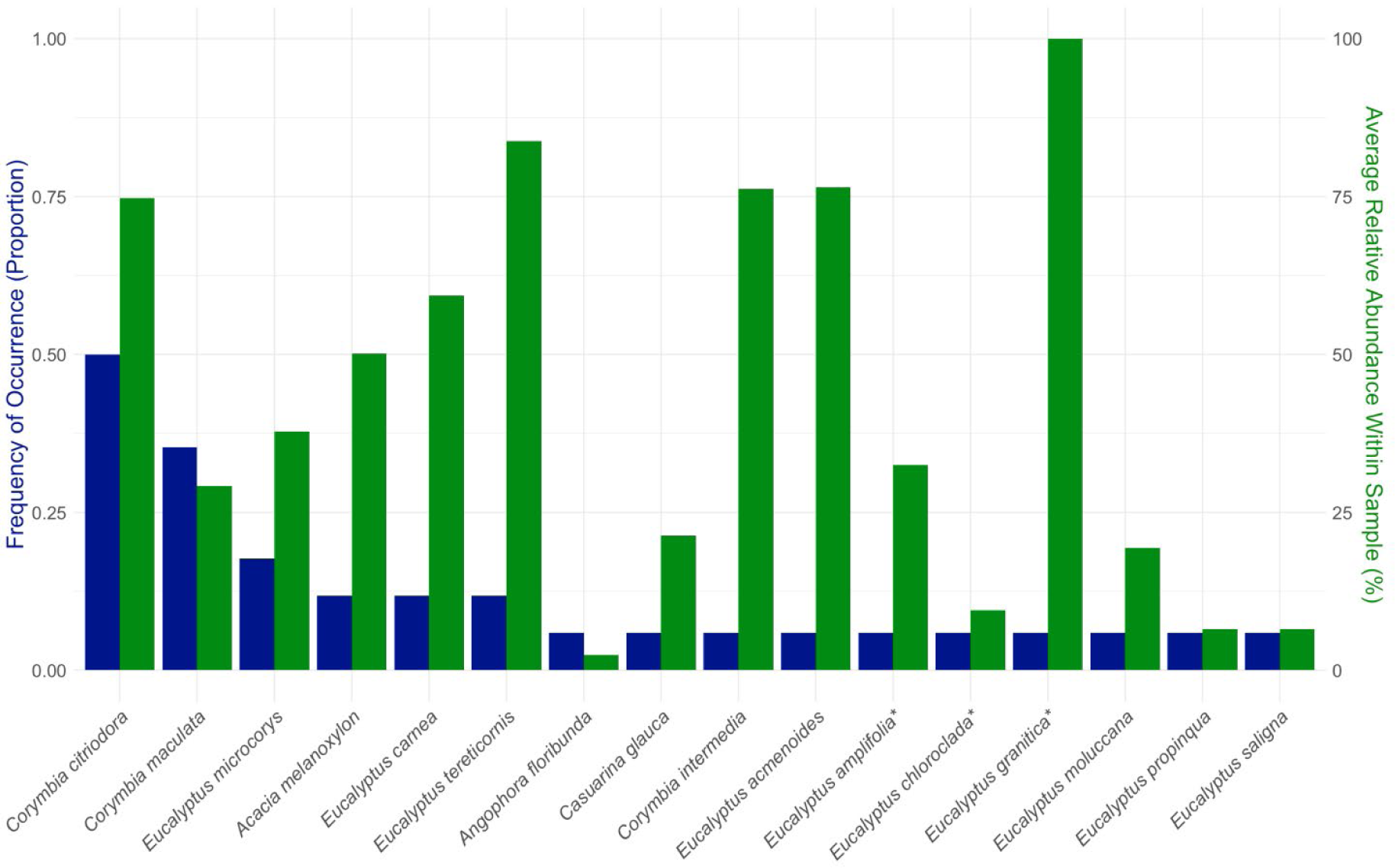
The frequency of occurrence and average relative abundance of plant species detected in the greater glider faecal samples. Left y-axis (blue) indicates the proportion of total faecal samples containing the plant species. Right y-axis (green) shows the relative abundance of each plant species within each individual sample they were detected in. *Natural range of species does not overlap with study sites.

### Comparisons to local species lists to identify potential false detections

A comparison with local species lists and known species distribution highlighted four plant species detected in the greater glider diet by DarTag sequencing whose naturally occurring range does not overlap our study region. These species were *Eucalyptus amplifolia, E. chloroclada, E. grantica* and *E. laevopinea.* Notably, all these species were only detected in a single sample with only 2 SNPs amplifying out of a possible 9-17.

## Discussion

Here, we used targeted SNP genotyping from faecal samples to identify feed tree species of greater gliders in two areas of southeast Queensland, Australia. Diet associated reads were detected in 71% of samples, demonstrating that the QLD Diet Oligo DArTag Panel designed for koalas can successfully identify plant species from greater glider faecal samples. Although, the higher failure rate and often lower read counts returned from the greater gliders compared to koalas (Blyton et al. 2023) suggests that not all dietary species were captured in the existing panel. We found a total of 17 plant species (13 supported by overlap of their distribution with the study sites) across five genera, with three species occurring across a range of sample locations and the remaining species being locally specific. This rejects our hypothesis that greater gliders feed on the same two or three species regardless of location, as most of the plant species were unique to the sampling locations. In general, this also runs contrary to the previous assumptions that greater gliders are highly specialised feeders, only consuming *Eucalyptus spp.,* as it appears their diet includes a number of local tree species across multiple genera that vary both between, and even within, sites. It should however be noted that the number of species identified was still relatively small, and the majority were from closely related genera. This runs consistent with the findings from koala DArTag dietary analysis (Blyton et al., 2023). From the species we identified, only *C. citriodora, E. tereticornis, C. intermedia, E. acmenoides* and *E. moluccana* have previously been documented as feed preferences of greater gliders through observational studies (Comport et al., 1996; Kehl & borsboom, 1984; Smith et al., 2007; Starr et al., 2021). Prior to our study, there were no recordings of greater gliders feeding on *Acacia, Angophora* or *Casuarina* trees. This suggests that the use of targeted genetic sequencing for this folivore diet provides a more extensive insight into plants being ingested by our target species compared to previous observational research methods.

In our study, we found variation in greater glider feed tree preferences between our sampling locations. We found similarities in species preferences between the Sheep Station Creek, Cedar Creek, Clear Mountain and Cashmere populations, however the Mount Glorious and Gympie gliders were feeding on distinctly different species. A factor that could have influenced the this is the availability of trees to feed on at each sample site. Without incorporating an aspect of thorough vegetation surveys into the project, we are unable to deduce if the greater gliders are feeding on certain species out of preference, or whether they are just eating what is available to them within their habitat. Koalas, for example, have been shown to rapidly change food preference to new eucalypt species after being translocated or provided with a plant they had previously not been exposed to (Blyton et al., 2023; Moore & Foley, 2000). We found that the greater gliders from Gympie were eating different species than those in Moreton Bay. This could be due to the Gympie sampling location having a different selection of feed tree species available compared to the Moreton Bay populations, which showed fairly consistent intra-location results. Although it was not explicitly recorded in our study, soil types and topography between the lower-lying Gympie region and the hillier areas of Moreton Bay, such as Mount Glorious and Clear Mountain, would vary greatly, cultivating a different composition of plant species (Cochrane et al., 2023; Singh et al., 2021).

We are unable to deduce the impact this had on our study, however we can hypothesise that tree species availability may have influenced the distinct feeding preferences seen between the Moreton Bay and Gympie samples. Alternatively, differences in the dietary preferences of the Gympie greater gliders may be due to them being a different species to those further south. The delineation between greater glider species remains poorly understood, particularly with their ability to hybridise (McGregor et al., 2020). Without further genetic testing of the individuals the samples were taken from, we are unable to determine if this may have impacted the dietary choices in our study.

Another possible factor contributing to the variation in greater glider feed tree preferences between our sampling locations is the presence or absence of competing arboreal based on the demand of ‘nutritional niche partitioning’ (Jensen et al., 2014). Previous research into nutritional aspects of arboreal folivore diet found that most species tend to find balance between the reward of higher nutritional intake without the detriment of too many toxins (Moore et al., 2004). Greater gliders can withstand higher levels of secondary metabolites compared to most other folivore species, particularly sideroxylonals (Jensen et al., 2014; Youngentob et al., 2011). This allows for ‘nutritional niche partitioning’ between greater gliders, koalas (*Phascolarctos cinereus*), brushtail possums (*Trichosurus vulpecula)* and ringtail possums (*Pseudocheirus peregrinus*) which often share overlapping home ranges (Jensen et al., 2014). The presence or absence of competing arboreal species may have influenced the dietary choices of the greater gliders in our study, accounting for the difference in diet across locations. For example, we found greater gliders feeding on *E. tereticornis* and *E. acmenoides* in the Mount Glorious and Gympie populations, both of which are species high in sideroxylonals, despite the presence of species lower in sideroxylonals (Supplementary Table 2). On the contrary, greater gliders in Sheep Station Creek and Clear Mountain were found to eat more *Corymbia* species that are lower in sideroxylonals. It is possible that there is more competition for food at Mount Glorious and Gympie, forcing the gliders to feed on species with the higher secondary metabolites and lower nitrogen levels, whereas the greater gliders with less competition can preference species with higher nutritional value and lower secondary metabolites. Future studies combining dietary analysis with arboreal fauna surveys could provide better insight into the impact on competition pressure and niche partitioning on dietary decision making of greater gliders.

To help optimise conservation efforts, future research should look at coupling dietary analysis with vegetation surveys to clarify which species are available to feed on, and which are preferred when choices are available. Furthermore, it would be valuable to explore whether different populations or individual gliders do favour specific feed tree species, even if the same selection of species was available to them. This has been seen in koalas on a population level, where *E. radiata* was untouched by koala populations in Walkerville, Victoria, even when they were suffering a loss of condition and increased mortality due to their preferred species being defoliated, yet *E. radiata* was a species of choice for koalas in northeastern New South Wales (Krockenberger, 1993; Martin, 1985). More recent research has discovered that differences in gastrointestinal microbiome between koala populations is a strong driver of their diet selection (Blyton et al., 2019). At an individual level, nothing is documented about how characteristics like sex, age class or reproductive demands alter the dietary needs of greater gliders. It is recognised that the diet of other arboreal folivores shift based on these factors, therefore it is possible that greater gliders would also be impacted. For example, female koalas have been shown to increase their food intake by 35% to meet the demands of reproduction (Krockenberger & Hume, 2007), and a direct correlation has been seen between nitrogen intake and reproductive success in common brushtail possums (DeGabriel et al., 2009). We are unable to say whether this impacted our findings due to a lack of information on the individuals our samples came from, therefore future research into the effects of population and individual differences will be important for a thorough understanding of the influence this has on greater glider diet.

Although they were only identified at low relative abundance, our study is the first to identify *Acacia* and *Casuarina* as feed trees for greater gliders. These findings hold promise for conservation and restoration as *Acacia* and *Casuarina* plants are known as pioneer species, often fast growing, even in unideal environments (Martínez-Ramos et al., 2021). Compared to eucalypts which take longer to grow (Taylor et al., 2014), planting *Acacia* or *Casuarina spp*. may offer a faster solution to bolster food availability in greater glider habitat and act as a supplementary food source as larger eucalypt trees establish. *Casuarina glauca* and *Acacia melanoxylon* were the only species from their respective genera included in the DarTag panel. Expanding the panel to include markers for additional *Casuarina* and *Acacia* species will be beneficial for identifying whether greater gliders are selective with the species they consume, or if a variety of species from these genera can be included in mixed species plantings for restoration projects.

Although we found the targeted species-specific sequencing to be successful at identifying feed tree preferences for greater gliders, the methodology does have limitations (Blyton et al., 2023). Targeted SNP sequencing is advantageous over other dietary analysis methods as it eliminates the chance of human error when visually identifying food sources, either as they are being ingested or in faecal pellets (Holechek et al., 1982). Additionally, it drastically reduces time spend in the field conducting focal surveys and spotlighting (Blyton et al., 2023; Gilby et al., 2010). However, the SNP sequencing is limited to the number of plant species in the reference panel, therefore increasing research into expanding the panel will produce a more comprehensive picture of dietary choices for greater gliders, koalas, and other folivores that can benefit from this analysis (Blyton et al., 2023). Similarly with DNA metabarcoding, the panel is subject to quantitative error as the digestibility of leaves varies among species (Garnick et al., 2018). This is of particular concern for greater gliders who are known to feed on young leaves, which often have a faster digestive time (Jensen et al., 2014).

A notable concerning finding from this study was the detection of four species in the greater gliders’ diet outside there natural range; *Eucalyptus amplifolia, E. chloroclada, E. grantica* and *E. laevopinea.* One potential reason for this discrepancy may be that the species have been planted outside their natural range. Alternatively, they could represent erroneous results due to off target amplifications. It is possible that the genomic sequences for these species closely resemble other plant species that were not included in the DarTag panel. Without testing the specificity of each marker against every possible food species at each study site, there is the possibility that they will amplify from species not represented in the panel. Due to the similarities of *Eucalyptus* genomes, it is likely that these species were detected in the place of other locally occurring *Eucalyptus spp*. that are not currently in the panel. This again highlights the importance of continuing to expand on the existing DarTag panel, increasing the overall accuracy of this methodology for future use.

This study found that *C. citriodora* and *C. maculata* are major components of the diet of greater gliders in Moreton Bay. Additionally, we show that greater gliders are not strictly eucalypt specialists but feed on *Acacia, Casuarina* and *Angophora* species, as well as the previously known *Eucalyptus* and *Corymbia*. Conservation plans considering these findings can be coupled with the existing landscape management plans to comprehensively protect all components of the habitat that are critical for greater glider survival. Moreover, this targeted species-specific SNP sequencing approach can be used as a framework for dietary analysis of other species beyond koalas and greater gliders, particularly those with a cryptic nature, although more work is needed to develop oligo panels that include all potential food species for each folivore and into developing corrections for differential digestibility. The information gained from these analyses can be used as a reference for implementing habitat restoration plans to conserve endangered species on a local and global scale.

## Author contributions

The project was conceived by D.A. Potvin, C.H. Frère and M. J. Bratovic. Fieldwork was completed by M.J. Bratovic. Laboratory work was completed by M.J. Bratovic with guidance by N. Jackson and L. Moss. Host DNA sequences analysed by N. Jackson. Data analysis was lead by M.J. Bratovic and supported by D.A. Potvin and M.D.J. Blyton. The initial manuscript draft was written by M.J. Bratovic. All authors contributed to editing and final manuscript preparation.

## Data availability

All relevant data has been provided in the supplementary material.

## Acknowledgements

We would like to thank City of Moreton Bay and the University of the Sunshine Coast for providing funding towards this project.

