## Supplementary Table 1 for "The future is faeces: Using faecal genomic sequencing to understand dietary choices of an endangered arboreal marsupial"

**Locations for collection of greater glider faecal samples**

| SITE NAME | SITE CODE | LATITUDE | LONGITUDE |
|---|---|---|---|
| CASHMERE | C | -27.2916 | 152.8926 |
| CEDAR CREEK | CC | -27.3335 | 152.8209 |
| CLEAR MOUNTAIN | CM | -27.3307 | 152.9098 |
| GYMPIE | G | -26.0349 | 152.5139 |
| MOUNT GLORIOUS | MG | -27.3333 | 152.7667 |
| SHEEP STATION CREEK | SSC | -27.1311 | 152.9000 |
